# Multimerization and tubulin binding are required for the SPIRAL2 protein to localize to and stabilize microtubule minus ends

**DOI:** 10.1101/2021.10.05.463238

**Authors:** Yuanwei Fan, Natasha Bilkey, Ram Dixit

## Abstract

Accruing evidence points to the control of microtubule minus-end dynamics as being crucial for the spatial arrangement and function of the microtubule cytoskeleton. In plants, the SPIRAL2 (SPR2) protein has emerged as a microtubule minus-end regulator that is structurally distinct from the animal minus-end regulators. Previously, SPR2 was shown to autonomously localize to microtubule minus ends and decrease their depolymerization rate. Here, we used *in vitro* and *in planta* experiments to identify the structural determinants required for SPR2 to recognize and stabilize microtubule minus ends. We show that SPR2 contains a single N-terminal TOG domain that binds to soluble tubulin. The TOG domain, a basic region, and coiled-coil domain are necessary and sufficient to target and stabilize microtubule minus ends. We demonstrate that the coiled-coil domain mediates multimerization of SPR2 that provides avidity for microtubule binding and is essential for binding to soluble tubulin. While TOG domain-containing proteins are traditionally thought to function as microtubule plus-end regulators, our results reveal that nature has repurposed the TOG domain of SPR2 to regulate microtubule minus ends.

## INTRODUCTION

Microtubules are cytoskeletal filaments composed of αβ-tubulin heterodimers that polymerize into linear protofilaments which associate laterally to form a hollow tube-like structure. Individual microtubules alternate randomly between periods of polymerization and depolymerization in a non-equilibrium process called dynamic instability (Mitchison and Kirschner, 1984). The two ends of a microtubule differ in their polymerization kinetics: the fast-growing end of a microtubule is called the plus end, while the slow-growing end is called the minus end. Cells regulate microtubule dynamics spatially and temporally to construct morphologically different arrays needed to accomplish vital tasks such as cell morphogenesis, cell cycle progression and intracellular transport.

Microtubule ends are particularly important for the regulation of microtubule dynamics because polymer growth and shortening occurs by the addition and removal of tubulin subunits at the ends. In cells, the growth and dynamics of microtubules occurs primarily at the microtubule plus ends (Desai and Mitchison, 1997). Traditionally, microtubule minus ends were considered to be quiescent in cells, but recent work has revealed otherwise and underscored the importance of minus-end dynamics to microtubule organization and stability (Akhmanova and Steinmetz, 2019). The dynamics of both microtubule ends are controlled by a group of specialized proteins which recognize specific tubulin conformations at the microtubule plus end and minus end, respectively (Akhmanova and Steinmetz, 2015; Nogales and Zhang, 2016). The most extensively studied microtubule plus-end regulatory proteins are the microtubule plus-end tracking proteins (+TIPs). These proteins accumulate specifically at microtubule plus ends during the growth phase and serve mainly to stabilize the plus end and to link it to other cellular structures (Galjart, 2010). Likewise, a group of proteins that specifically localize to free microtubule minus ends are called microtubule minus-end targeting proteins (−TIPs). While −TIPs are structurally distinct from +TIPs, they perform similar functions in that they mainly stabilize free microtubule minus ends and attach minus ends to other cellular structures (Akhmanova and Steinmetz, 2019).

In animals, the calmodulin-regulated spectrin-associated protein (CAMSAP)/Patronin/Nezha family of −TIPs has been studied extensively. Different members of this protein family either track growing microtubule minus ends (Jiang et al., 2014), progressively accumulate on the microtubule lattice formed by growth at the minus end (Jiang et al., 2014), or bind to and inhibit the growth of minus ends (Hendershott and Vale, 2014). This protein family is distinguished by the presence of a C-terminal CAMSAP1, KIAA1078 and KIAA1543 (CKK) domain which confers microtubule minus-end targeting (Atherton et al., 2017; Atherton et al., 2019).

Plants do not possess homologs of the CAMSAP/Patronin/Nezha protein family and recently it was shown that the plant-specific SPIRAL2 (SPR2) protein autonomously localizes to microtubule minus ends and stabilizes them (Fan et al., 2018; Leong et al., 2018; Nakamura et al., 2018). The frequency and rate of depolymerization of microtubule minus ends are greatly enhanced in *spr2* mutants compared to wild-type plants (Fan et al., 2018; Leong et al., 2018; Nakamura et al., 2018). Notably, the SPR2 protein differs from the CAMSAP/Patronin/Nezha protein family in both structure and dynamics. SPR2 lacks the CKK domain that is required by the CAMSAP/Patronin/Nezha proteins to recognize microtubule minus ends. In addition, unlike the CAMSAP/Patronin/Nezha proteins, SPR2 tracks depolymerizing microtubule minus ends, localizes to the microtubule lattice, and to a lesser extent to growing microtubule plus ends (Fan et al., 2018; Nakamura et al., 2018).

The SPR2 protein contains five predicted Huntington, elongation factor 3, phosphatase 2A, target of rapamycin 1 (HEAT) repeats at the amino terminus followed by a basic region and coiled-coil domain (Fan et al., 2018). This domain architecture resembles the tumor overexpressed gene (TOG) domain-containing microtubule regulatory proteins XMAP215 and CLASP, which localize primarily to microtubule plus ends. TOG domains are characterized by the presence of six HEAT repeats that form a paddle-shaped structure that binds with high affinity to a tubulin dimer (Al-Bassam et al., 2007; Slep and Vale, 2007; Ayaz et al., 2012). While TOG domains are best known for their ability to bind to free tubulin subunits (Ayaz et al., 2012; Ayaz et al., 2014), structural changes in some of the TOG domains of XMAP215 proteins might alter their binding preference to tubulin that is part of the microtubule lattice (Fox et al., 2014). Proteins that contain more than two TOG domains (e.g., human and plant XMAP215 homologs) function as monomers (Currie et al., 2011; Lechner et al., 2012; Widlund et al., 2011), whereas those that contain only one or two TOG domains (e.g., yeast XMAP215 homolog and CLASP) function either as dimers or oligomers (Gard et al., 2004; Al-Bassam et al., 2006), with the possible exception for the mitogen-activated protein kinase activator MEKK1 (Filipcik et al., 2020). Thus, the presence of multiple TOG domains is an important characteristic of proteins that regulate microtubule dynamics.

In this study, we conducted structure-function analysis of the SPR2 protein to elucidate the structural determinants for recognizing and stabilizing microtubule minus ends. We found that both the N-terminal HEAT repeat domain and basic region are required for SPR2 to bind to microtubules and that multimerization mediated by the coiled-coil domain greatly increases the affinity of SPR2 for microtubules. By contrast, the C-terminal domain does not contribute to this activity. Furthermore, we demonstrate that multimerization of SPR2 is essential for the N-terminal domain to bind to free tubulin dimers. When expressed in the *Arabidopsis thaliana spr2-2* mutant, the N-terminal region and coiled-coil domain of SPR2 were both necessary and sufficient to stably bind to the minus ends of cortical microtubules and inhibit their depolymerization rate. Together, our results establish SPR2 as a TOG domain-containing protein that has evolved to regulate the dynamics of microtubule minus ends.

## RESULTS

### The N-terminal region of SPR2 mediates microtubule binding

SPR2 is a relatively large, multi-domain protein (Fan et al., 2018). To determine which domains allow SPR2 to bind to and stabilize microtubule minus ends, we generated various truncated versions of the SPR2 protein that were tagged with green fluorescent protein (GFP) at the C-terminus (Figure 1a). Full-length and truncated GFP-labeled SPR2 proteins were recombinantly expressed and purified (Figure 1b) and used to assess microtubule binding in functional reconstitution experiments. These *in vitro* experiments consist of dynamic rhodamine-labeled microtubules that are polymerized from immobilized GMPCPP-stabilized microtubule fragments (called “seeds”). We found that the N-terminal fragment of SPR2 that lacks the basic region (1-276 aa) does not bind to microtubules (Figure 2a). Inclusion of the basic region conferred weak binding to the microtubule lattice, with the extent of microtubule binding correlating positively with the length of the basic region (compare the 1-340 aa and 1-400 aa fragments in Figure 2a). The largest N-terminal fragment without the coiled-coil domain (1-500 aa) showed both microtubule lattice and minus end binding (Figure 2a).

**Figure 1.**
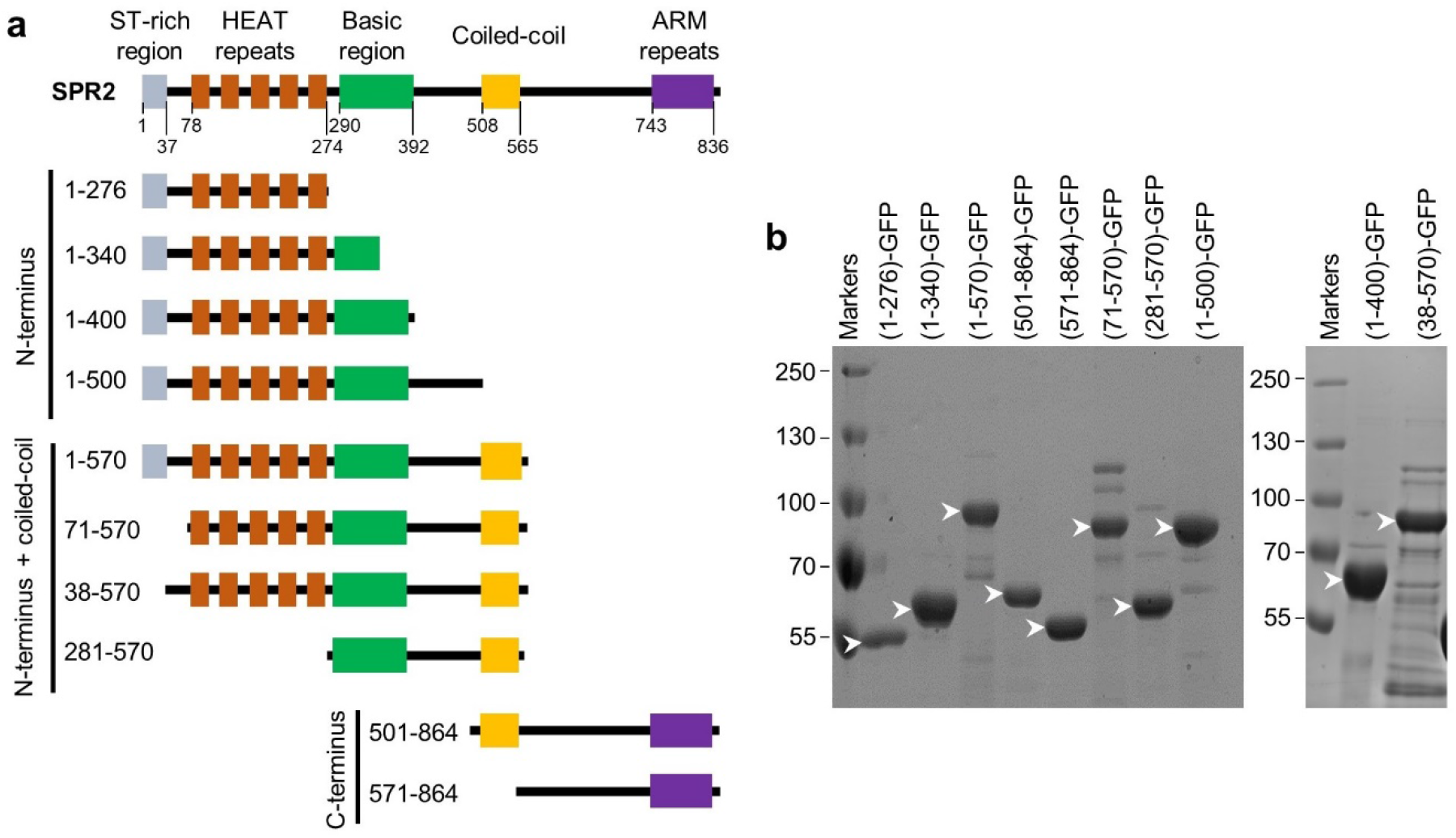
SPR2 domain organization. (**a**) Schematic representation of the domain architecture of the SPR2 protein and the various truncations used in this study. (**b**) Coomassie Blue-stained SDS-PAGE gels of recombinantly expressed and purified GFP-tagged SPR2 proteins used in this study. Arrowheads indicate the expected size of each protein.

**Figure 2.**
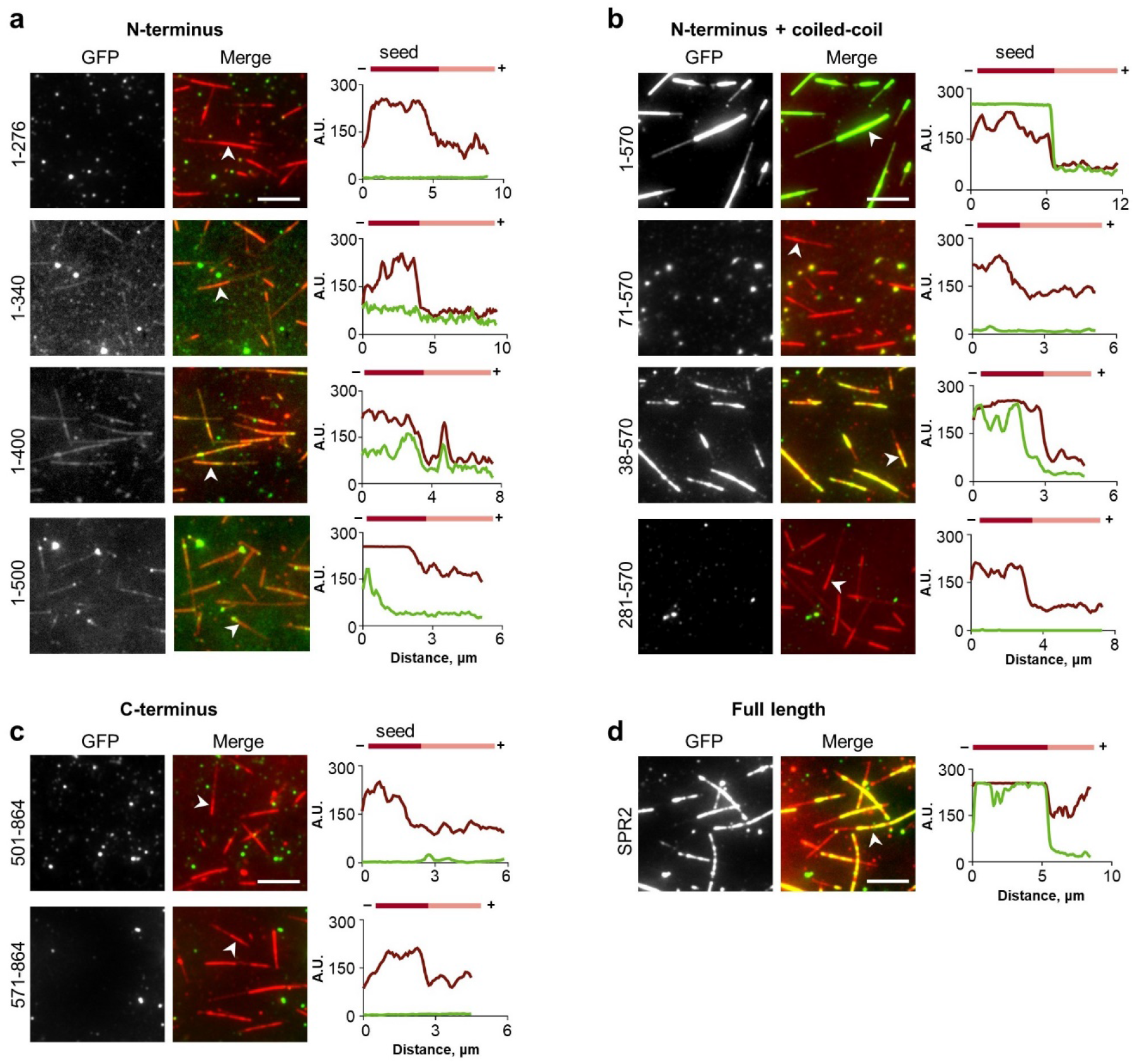
Analysis of microtubule binding and minus-end localization of the SPR2 fragments. Representative images of GFP-tagged N-terminal fragments (**a**), N-terminus and coiled-coil domain containing fragments (**b**), C-terminal fragments (**c**), and full-length SPR2 (**d**). Micrographs of the GFP channel alone and merge of the red rhodamine-labeled tubulin channel and green GFP channel are shown. Arrowheads label the microtubules whose fluorescence intensity profiles are shown. A.U., arbitrary units. Scale bar = 5 μm.

The coiled-coil domain dramatically increased the microtubule binding ability of the N-terminal fragment of SPR2 (1-570 aa). The 1-570 aa fragment bound more readily to the GMPCPP-stabilized microtubule seeds than to the GTP/GDP-state dynamic polymer (Figure 2b). This binding pattern is similar that of the full-length SPR2 protein (Fan et al., 2018). The strong microtubule binding of the 1-570 aa fragment was abrogated by deleting the first 70 amino acids (71-570 aa, Figure 2b). However, strong microtubule binding is restored when only the first 37 amino acids are deleted (38-570 aa, Figure 2b). Interestingly, the SPR2 fragment spanning just the basic region and coiled-coil domain (281-570 aa) is unable to bind to microtubules (Figure 2b). Similarly, the C-terminal region of the SPR2 protein, with or without the coiled-coil domain, does not bind to microtubules (Figure 2c).

Comparison of the microtubule binding of the various SPR2 protein fragments to the full-length SPR2 protein (Figure 2d) indicates that the 38-570 aa fragment most closely mimics the extent of microtubule binding and pattern of microtubule localization of the full-length SPR2 protein.

### Multimerization of SPR2 enhances microtubule binding

Given the striking difference in the extent of microtubule decoration between the 1-500 aa and 1-570 aa fragments of SPR2, we investigated whether the coiled-coil domain increases the microtubule binding affinity of SPR2. Using microtubule co-sedimentation assays, we found that the 1-500 aa fragment shows nonsaturating microtubule binding with a KD of 17.4 μM (Figure 3a and 3c). By contrast, the 1-570 aa fragment bound to microtubules with a KD of 1.3 μM (Figures 3b and 3d). Therefore, the coiled-coil domain increases the microtubule binding affinity of SPR2 by at least one order of magnitude.

**Figure 3.**
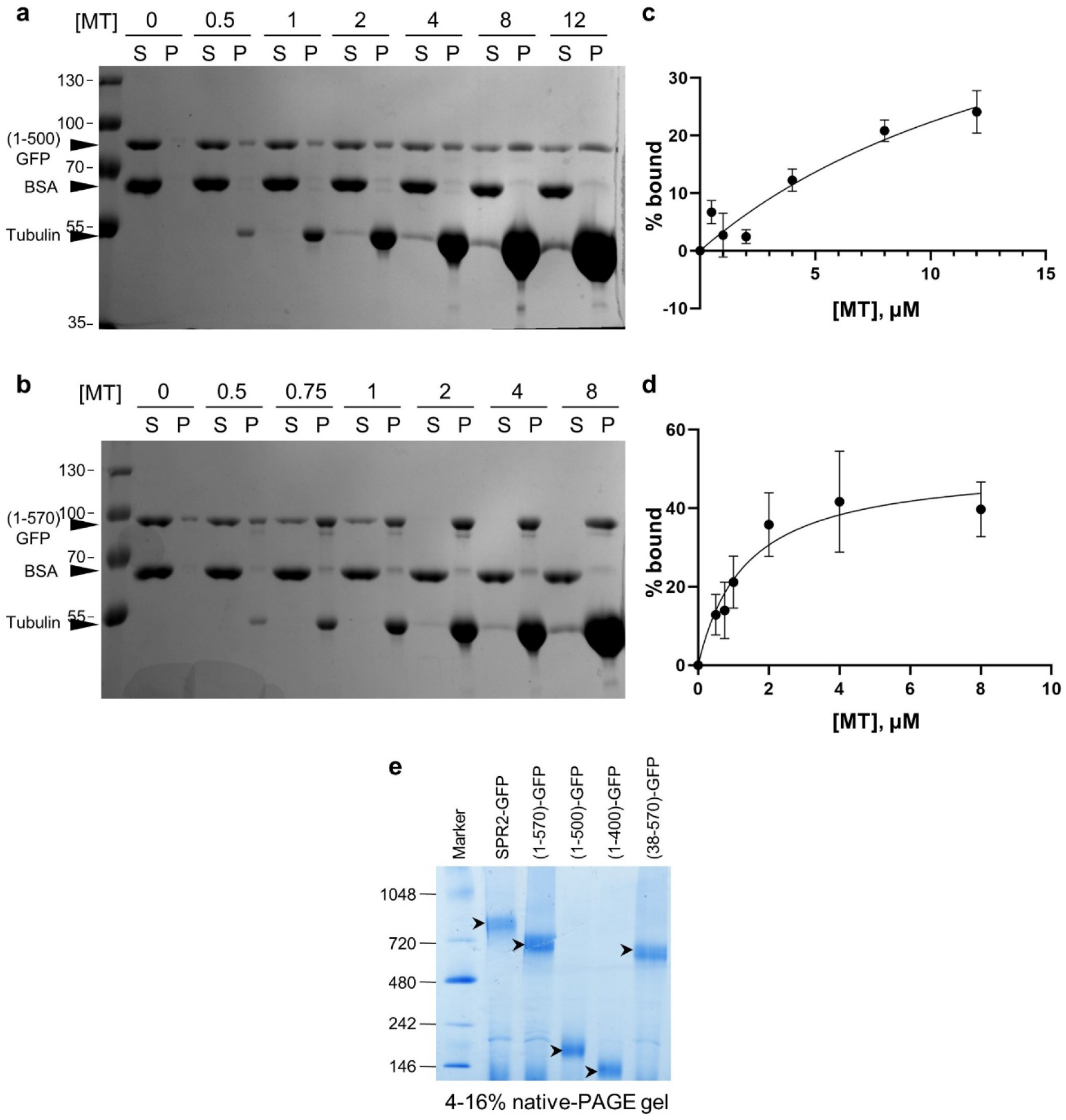
Multimerization state of SPR2 fragments. (**a**, **b**) Coomassie Blue-stained SDS–PAGE gels of microtubule cosedimentation assays. 2 μM (1-500)-GFP (**a**) or (1-570)-GFP (**b**) were coincubated with increasing concentrations of taxol-stabilized microtubules, 0.1 mg/ml bovine serum albumin (BSA), and 20 μM taxol for 20 min. S, supernatant; P, pellet. Arrowheads identify the different protein bands. (**c**, **d**) Microtubule binding curves for (1-500)-GFP (**c**) or (1-570)-GFP (**d**), respectively. Each data point represents the mean ± SD from three independent experiments. The data were fit to a model assuming single-site specific binding yielding KD values of 17. 4 μM and 1.3 μM for (1-500)-GFP and (1-570)-GFP, respectively. (**e**) NativePAGE 4-16% Bis-Tris gel of full-length SPR2 protein and indicated SPR2 fragments.

Yeast two-hybrid experiments have demonstrated that SPR2 interacts with itself (Fan et al., 2018). In addition, photobleaching analysis indicates that SPR2 exists primarily as multimers (Fan et al., 2018). To determine whether the coiled-coil domain is responsible for multimerization of SPR2, we conducted native polyacrylamide gel electrophoresis of purified full-length SPR2-GFP protein and the 1-400 aa, 1-500 aa, 1-570 aa and 38-570 aa SPR2 fragments. We found that the SPR2 proteins containing the coiled-coil domain form higher-order multimers, whereas the SPR2 proteins without the coiled-coil domain did not (Figure 3e). Taken together, these data indicate that the coiled-coil domain drives multimerization of SPR2 that provides avidity for microtubule binding.

### The 1-500 aa, 1-570 aa and 38-570 aa fragments of SPR2 inhibit the dynamics of microtubule minus ends

Our functional reconstitution experiments showed that the 1-500 aa, 1-570 aa and 38-570 aa fragments of SPR2 localize to the minus ends of GMPCPP-stabilized microtubules (Figure 4a). To determine whether these SPR2 fragments affect the dynamics of microtubule plus and minus ends, we conducted kymograph analysis (Figure 4b). We included the 1-400 aa fragment in this analysis to assess whether its weak binding to microtubules influences microtubule dynamics.

**Figure 4.**
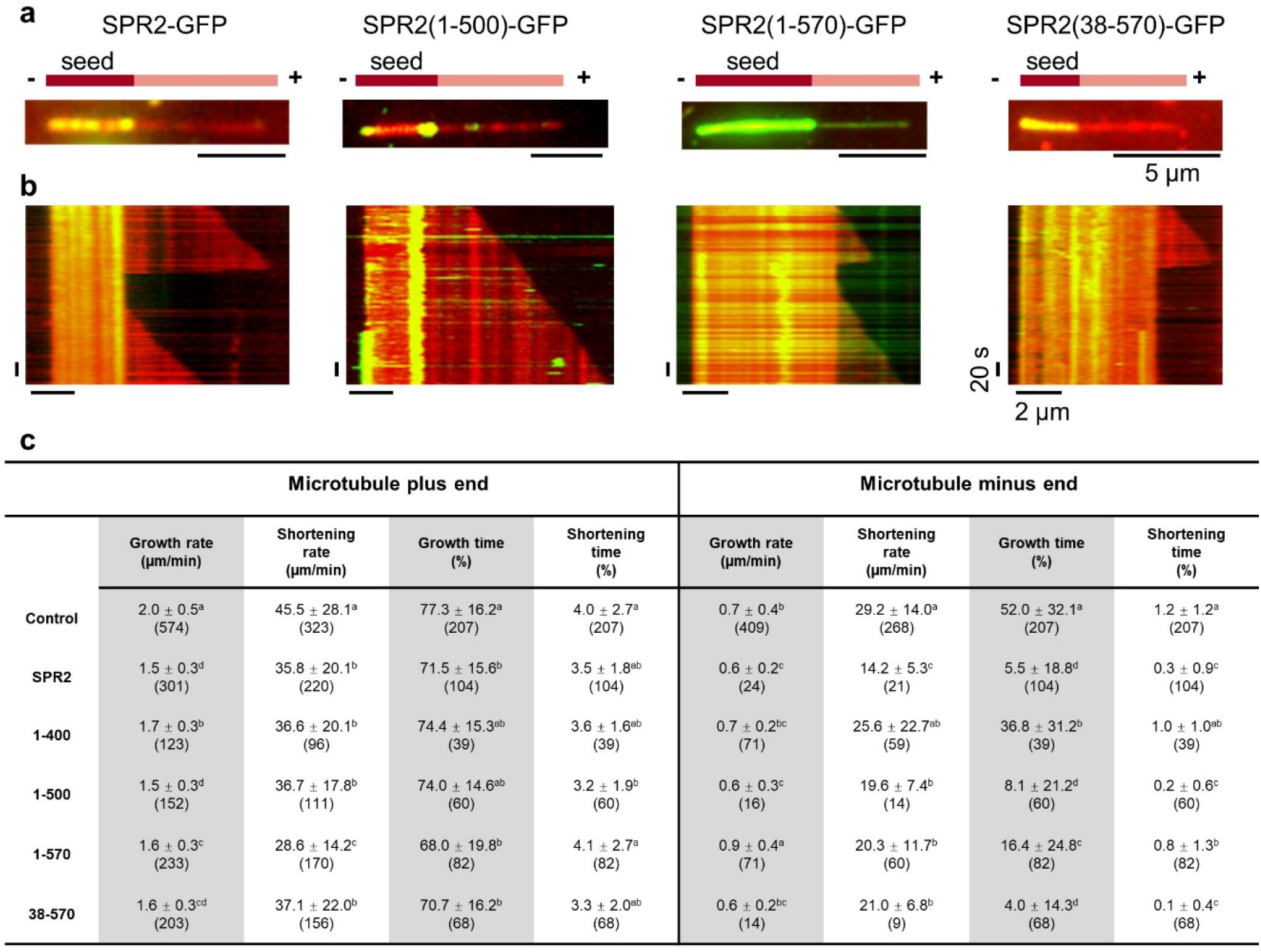
Effect of SPR2 fragments on microtubule dynamics *in vitro*. (**a**) Images showing the localization of 500 nM SPR2-GFP, (1-500)-GFP, (1-570)-GFP, and (38-570)-GFP proteins (green) on rhodamine-labeled dynamic microtubules assembled from GMPCPP-stabilized microtubule seeds (red). Scale bar = 5 μm. (**b**) Kymographs of the microtubules shown in (**a**). (**c**) *In vitro* microtubule dynamics measurements for the experiments shown in (**a**). Only microtubules that had continuous GFP signal at the minus end were analyzed. Values are means ± SD (n = number of events for growth and shortening rates; n = total number of kymographs for percent growth and shortening times). The letters indicate statistically distinguishable groups as determined by the Student’s *t* test (p < 0.05).

Microtubule minus ends that had a clearly detectable GFP signal from the 1-500 aa, 1-570 aa and 38-570 aa fragments spent significantly less time growing and shortening compared to the tubulin-alone control (Figure 4c). This effect is similar to that of the full-length SPR2 protein. Unexpectedly, the 1-570 aa fragment, which showed the highest microtubule binding, inhibited microtubule minus end dynamics to a lesser degree than the 1-500 aa and 38-570 aa fragments (Figure 4c).

The 1-400 aa fragment impacted microtubule minus end dynamics significantly less than the 1-500 aa, 1-570 aa and 38-570 aa SPR2 fragments (Figure 4c). All the SPR2 fragments used in this study altered the dynamics of microtubule plus ends to a small degree (Figure 4c). This is likely an indirect effect of these proteins binding to the microtubule lattice, as previously reported for full-length SPR2 (Fan et al., 2018).

### SPR2 tracks both growing and shortening microtubule minus ends *in vitro*

During our *in vitro* experiments, we found that the 1-570 aa fragment of SPR2 frequently tracks microtubule minus ends during both the growth and shortening phases (Figure 5a and 5b). This behavior is distinct from the CAMSAP/Patronin/Nezha protein family, members of which track microtubule minus ends only during the growth phase (Jiang et al., 2014). While we have previously reported that full-length SPR2 tracks shortening microtubule minus ends *in vitro* (Fan et al, 2018), here we show that full-length SPR2 also tracks growing microtubule minus ends (Figure 5c and 5d). Similar to the full-length SPR2 protein, the 38-570 aa fragment binds predominantly stably to microtubule minus ends (Figure 5e). However, unlike the full-length SPR2 protein, the 38-570 aa fragment does not track either growing or shortening minus ends (Figure 5e). This finding suggests that the Ser/Thr-rich region at the extreme N-terminus of SPR2 somehow contributes to its ability to track dynamic microtubule minus ends *in vitro*.

**Figure 5.**
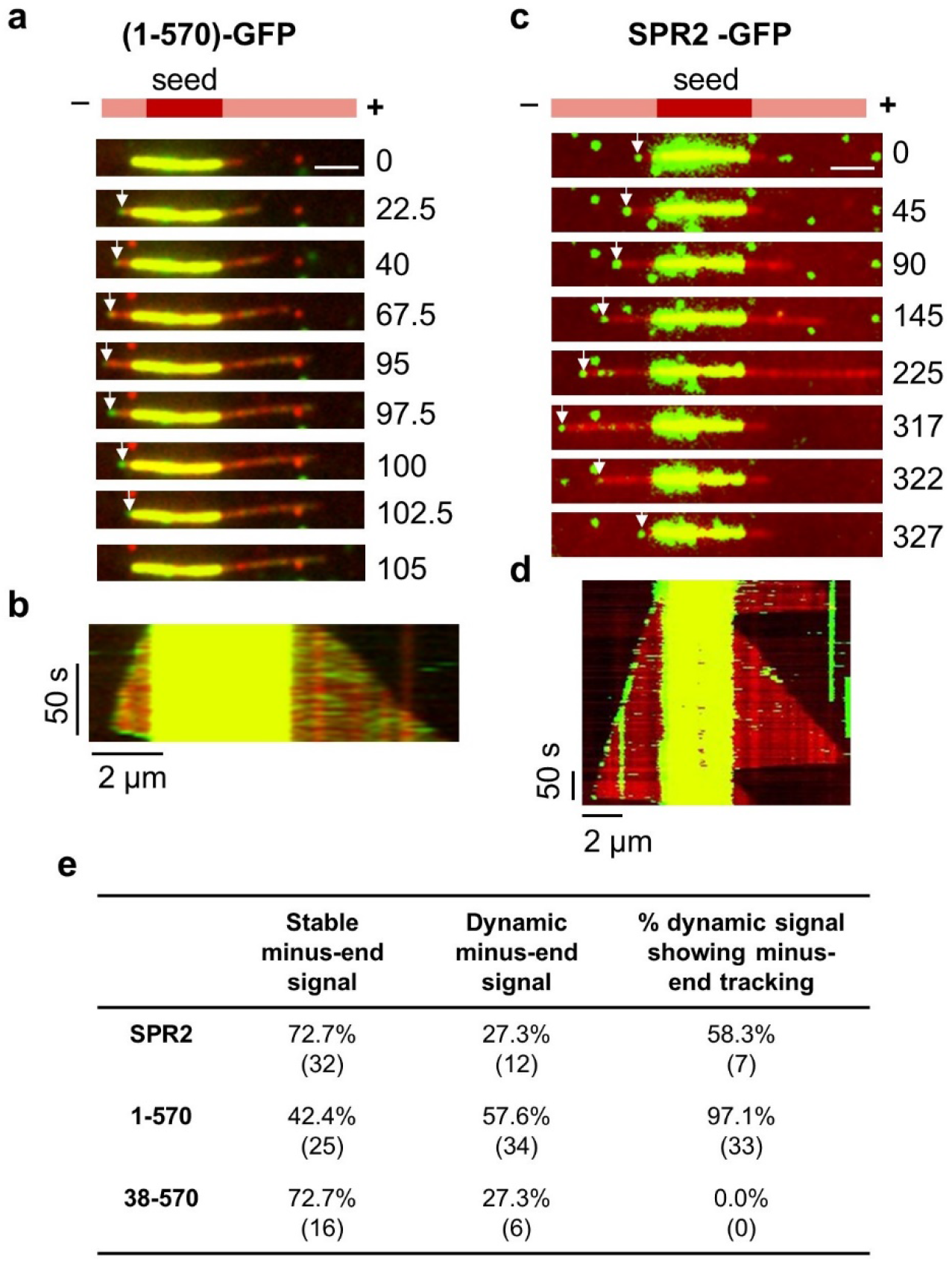
SPR2 tracks both growing and shortening microtubule minus ends. (**a**, **c**) Montage of the dynamics of plus and minus ends of rhodamine-labeled microtubules incubated with either 500 nM (1-570)-GFP (**a**) or 500 nM SPR2-GFP (**c**). Numbers show time in seconds. Arrows mark the minus end. Scale bar = 2 μm. (**b**, **d**) Kymographs of microtubules shown in (**a**) and (**c**), respectively. (**e**) Proportion of stable minus end binding and dynamic minus-end tracking by full-length SPR2-GFP, (1-570)-GFP, and (38-570)-GFP proteins.

### SPR2 binds to free tubulin dimers

The SPR2 protein is predicted to contain five HEAT repeats near the N-terminus (Figure 1a). We wondered whether SPR2 has a cryptic sixth HEAT repeat, which would constitute a TOG domain. Since TOG domains are best known for their ability to bind to free tubulin dimers (Ayaz et al., 2012; Ayaz et al., 2014), we conducted pulldown experiments to test this possibility with full-length SPR2-GFP. These experiments revealed that SPR2-GFP can bind directly to soluble tubulin (Figure 6a). To further analyze this activity and to determine which fragments of SPR2 bind to tubulin, we used a total internal reflection fluorescence (TIRF) microscopy-based assay. In this assay, unlabeled microtubules are co-incubated with a GFP-labeled SPR2 protein and rhodamine-labeled tubulin to determine whether microtubule-bound SPR2 proteins can simultaneously bind to free tubulin. In the absence of SPR2 protein, rhodamine-labeled tubulin did not localize to microtubules. However, in the presence of full-length SPR2-GFP protein, we observed punctate rhodamine-tubulin signal that coincides with SPR2-GFP puncta on the microtubule lattice (Figure 6b). Robust localization of rhodamine-tubulin to the microtubule lattice was observed with the 1-570 aa and 38-570 aa SPR2 fragments, but not with the 1-500 aa and 1-400 aa fragments (Figure 6b). Therefore, we conclude that multimerization of SPR2 is required to bind tubulin dimers.

**Figure 6.**
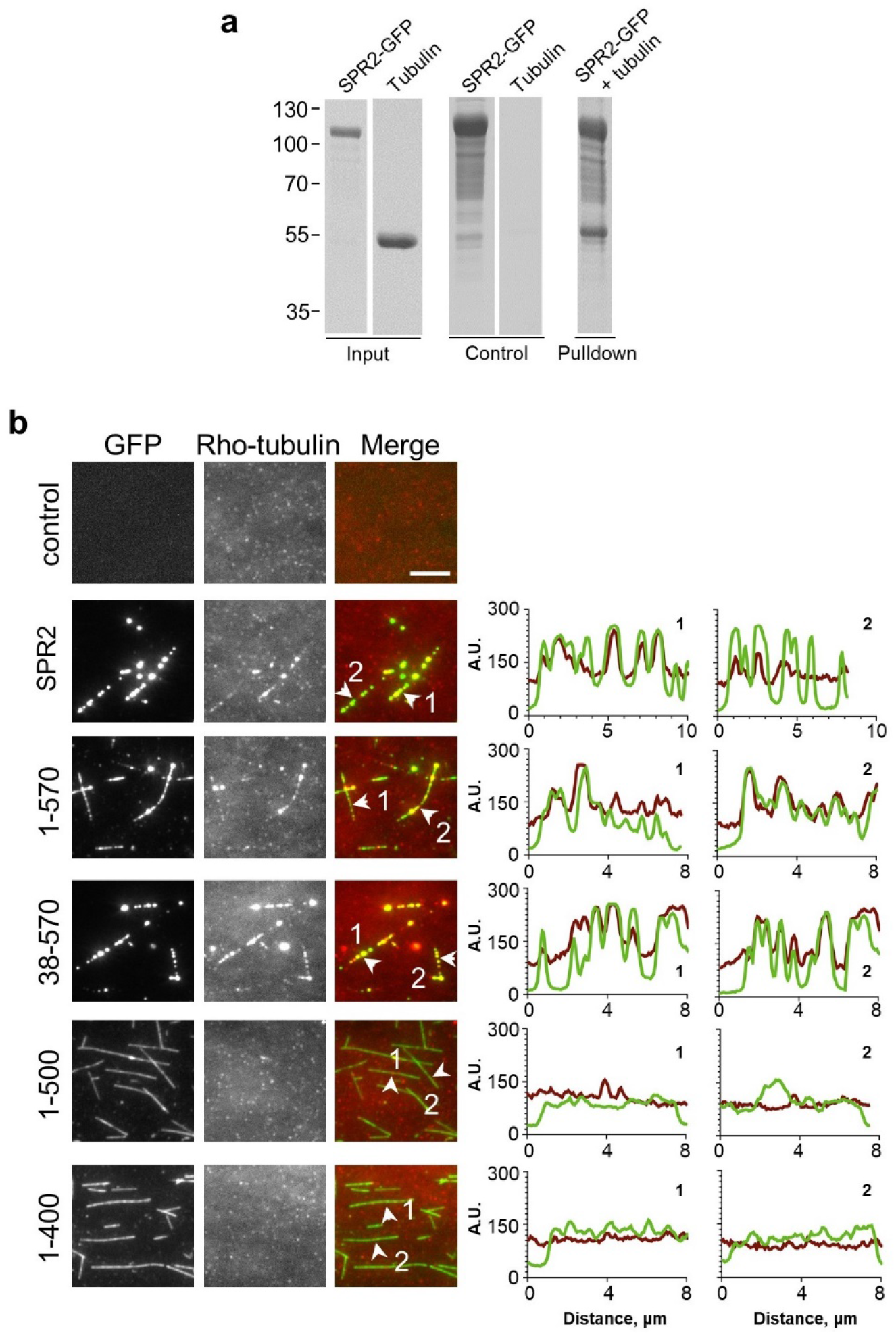
SPR2 binds soluble tubulin dimers. (**a**) Coomassie Blue-stained gel of an *in vitro* pulldown experiment with GFP-tagged full-length SPR2 incubated with soluble tubulin. (**b**) Micrographs of the indicated GFP-tagged SPR2 proteins incubated with unlabeled GMPCPP-stabilized microtubules and soluble rhodamine-labeled tubulin. Arrowheads and numbers label the microtubules whose fluorescence intensity profiles are shown. A.U., arbitrary units. Scale bar = 5 μm.

### The 1-570 aa and 38-570 aa fragments of SPR2 rescue the helical growth phenotype of the *spr2-2* mutant

To find out whether the SPR2 fragments that localize to and stabilize microtubule minus ends *in vitro* are functional *in planta*, we used genetic complementation of the *spr2-2* knockout mutant, which has a strong right-handed helical growth phenotype (Shoji et al., 2004).

The coding sequence of 1-500 aa, 1-570 aa and 38-570 aa fragments fused to mRuby were expressed under the control of the *SPR2* promoter in the *spr2-2* mutant containing a GFP-TUB6 microtubule marker. For the 1-500 aa fragment, we analyzed six independent single-insertion homozygous lines. All these plants were morphologically indistinguishable from the *spr2-2* mutant and exhibited right-handed helical growth of cotyledons, true leaves, and petals (Figure 7). For the 1-570 aa fragment, we analyzed eight independent single-insertion homozygous lines. Of these, two lines appeared like wild-type plants, indicating full complementation (e.g., line #2-4 in Figure 7). The remaining six (1-570)-mRuby expressing plants showed partial complementation. In these plants, the cotyledons still showed right-handed helical growth, but the true leaves and petals appeared like wild type (e.g., line #23-3 in Figure 7).

**Figure 7.**
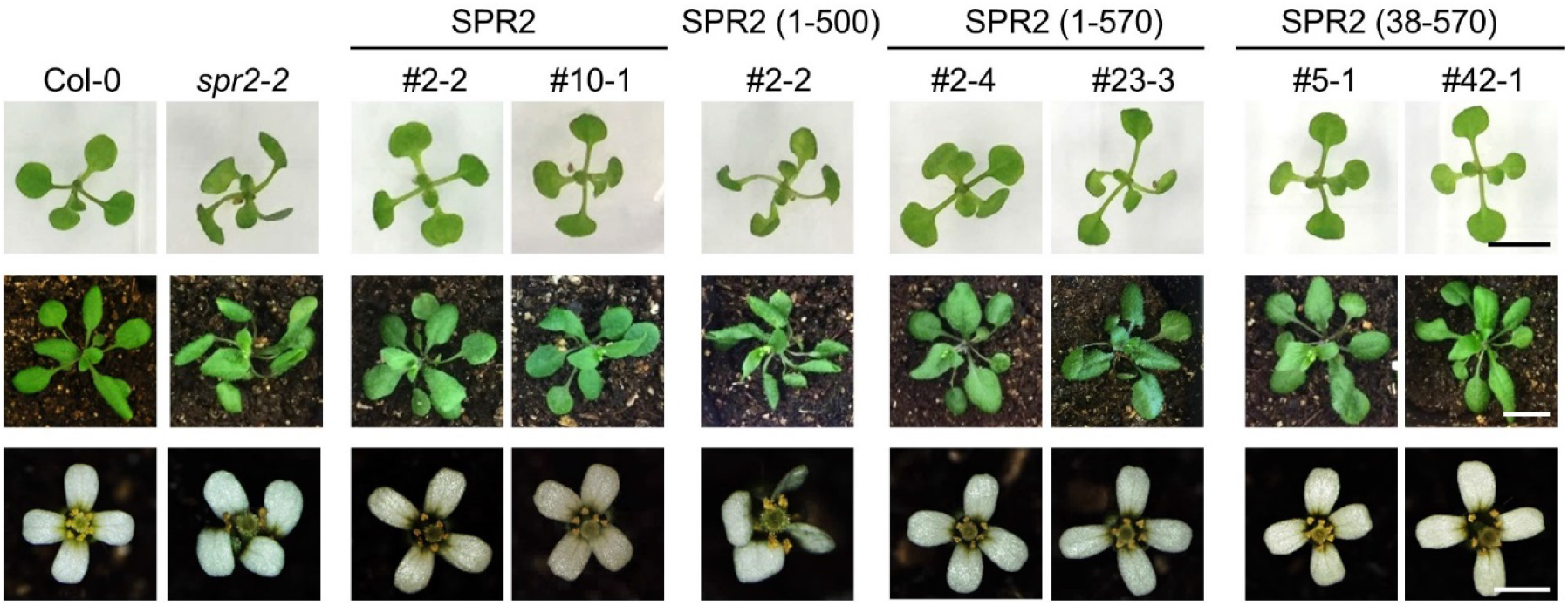
Phenotypes of the complemented *spr2-2* plants. Images of Col-0, *spr2-2* mutant, and *spr2-2* mutant complemented with either SPR2-mRuby, (1-500)-mRuby, (1-570)-mRuby, or (38-570)-mRuby fusion proteins. The top row shows seedlings 10 days after germination. The middle row shows 20-day-old plants grown in soil. The bottom row shows flowers from adult plants. Scale bar = 0.5 cm for seedling and plants; scale bar = 1 mm for flowers.

For the 38-570 aa fragment, we analyzed five independent single-insertion homozygous lines. Three of these lines showed full complementation, with cotyledons, true leaves, and petals showing wild-type-like morphology (e.g., lines #5-1 and #42-1 in Figure 7). The remaining two lines did not show any mRuby signal, probably because of gene silencing, and therefore were not analyzed further. As reported previously (Fan et al., 2018), expression of full-length SPR2-mRuby in the *spr2-2* mutant leads to complete complementation (Figure 7).

### The 1-570 aa and 38-570 aa fragments of SPR2 affect the dynamics of cortical microtubules *in planta*

We next performed live imaging of cortical microtubules to determine whether the genetic complementation results can be explained by the microtubule localization pattern of the different SPR2 fragments and by how they affect cortical microtubule dynamics.

The (1-500)-mRuby signal appeared only sporadically at the minus ends of cortical microtubules (Figure 8a) and these minus ends depolymerized at a fast rate (Figure 8b). In the 1-570 aa #2-4 line which shows full complementation, we found that the mRuby signal is occasionally absent from the minus end, which then exhibits fast depolymerization. However, once (1-570)-mRuby localizes to the minus end, the minus end becomes much more stable and depolymerizes slowly (Figure 8c and 8d). By contrast, in the 1-570 aa #23-3 line which shows only partial complementation, the mRuby signal at the cortical microtubule minus end is always discontinuous and the minus-end shortening rate and duration are intermediate compared to the *spr2-2* mutant and control plants (Figure 8e and 8f). Lastly, in the 38-570 aa complementation lines, the mRuby signal is present continuously at free microtubule minus ends (Figure 8g) which exhibit slow depolymerization (Figure 8h).

**Figure 8.**
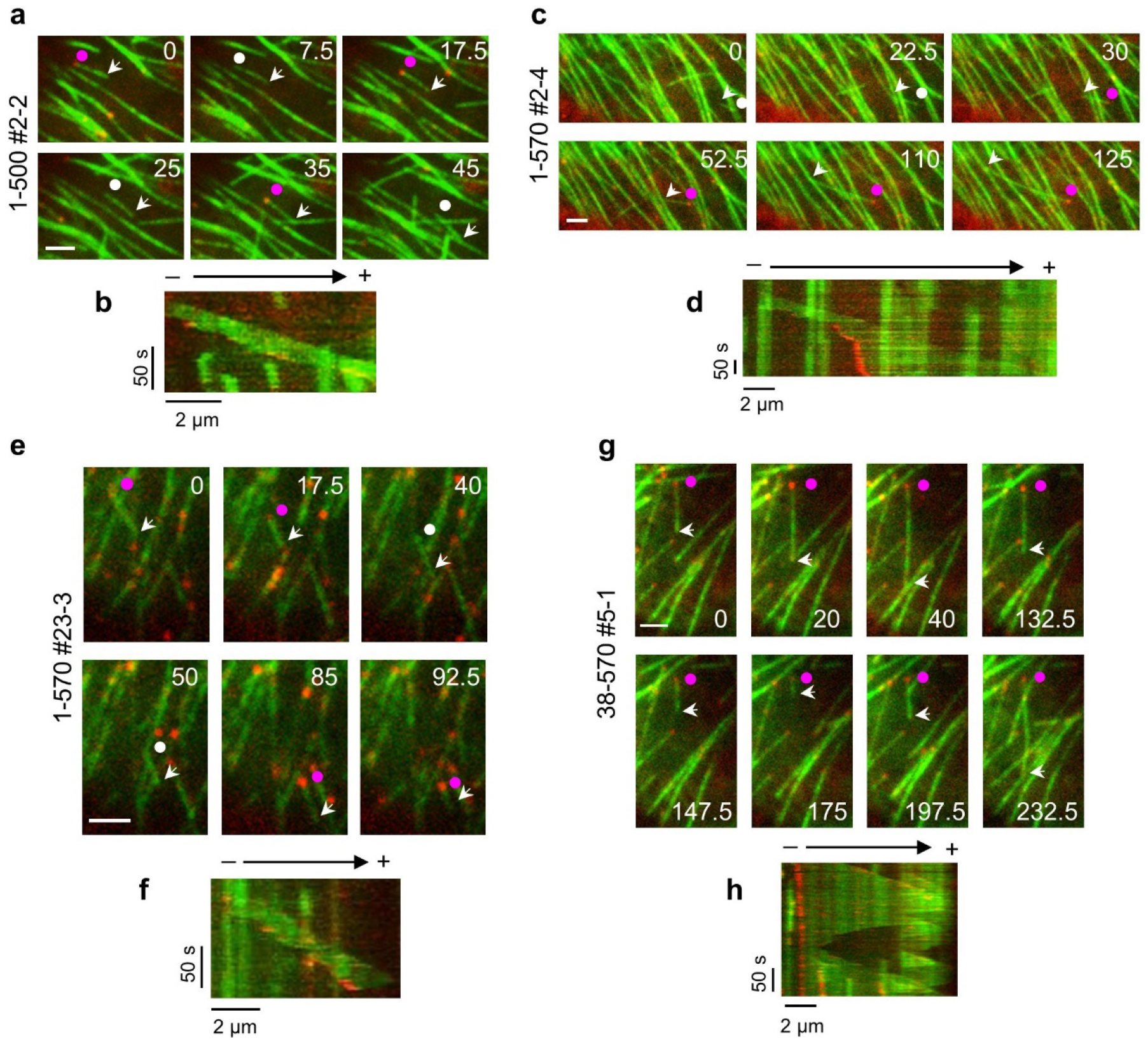
Cortical microtubule dynamics in complemented *spr2-2* plants. Montage and kymograph showing the localization and cortical microtubule dynamics of (1-500)-mRuby (**a**, **b**), (1-570)-mRuby line #2-4 that shows full complementation (**c**, **d**), (1-570)-mRuby line #23-3 that shows partial complementation (**e**, **f**), and (38-570)-mRuby line (**g**, **h**). Numbers show time in seconds. The arrow and circle label the microtubule plus and minus end, respectively. The purple circle indicates when the mRuby signal is present at the minus end. The white circle indicates when the mRuby signal is absent at the minus end. Scale bar = 2 μm.

We used these data to quantify the growth and shortening rates as well as the time spent growing, shortening, and pausing for both the plus and minus ends of cortical microtubules. The 1-500 aa, 1-570 aa and 38-570 aa fragments partially rescued the growth rate of the plus ends, except for the 1-570 aa line #23-3 in which the plus-end growth rate is statistically indistinguishable from control plants (Table 1). The microtubule plus-end shortening rate was statistically the same between all the lines (Table 1). For time spent by the plus ends in the various phases, the 1-500 aa lines failed to restore these parameters to control values; in fact, the deviation from control values was even more severe in these plants compared to the *spr2-2* mutant. By contrast, the 1-570 aa and 38-570 aa lines showed at least partial rescue of these parameters, with 38-570 aa line #5-1 being the most like control plants (Table 1).

**Table 1:** Dynamics of cortical microtubule plus and minus ends

| | | Growth rate<br>($\mu\text{m}/\text{min}$) | Shortening rate<br>($\mu\text{m}/\text{min}$) | Growth time<br>(%) | Shortening time<br>(%) | Pause time<br>(%) |
| --- | --- | --- | --- | --- | --- | --- |
| Microtubule plus end | SPR2 | $4.8 \pm 1.8^c$<br>(114) | $20.1 \pm 22.5^a$<br>(79) | $88.7 \pm 10.0^e$<br>(75) | $11.3 \pm 10.0^a$<br>(75) | 0<br>(75) |
| | *spr2-2* | $8.1 \pm 5.1^a$<br>(59) | $20.6 \pm 12.4^a$<br>(9) | $98.1 \pm 6.0^b$<br>(53) | $1.9 \pm 6.0^{de}$<br>(53) | 0<br>(53) |
| | 1-500<br>#2-2 | $6.2 \pm 2.3^b$<br>(62) | $11.4 \pm 5.3^a$<br>(2) | $99.9 \pm 1.2^a$<br>(61) | $0.2 \pm 1.2^e$<br>(61) | 0<br>(61) |
| | 1-570<br>#2-4 | $5.7 \pm 1.7^b$<br>(63) | $22.0 \pm 13.6^a$<br>(26) | $93.0 \pm 11.9^{cd}$<br>(51) | $7.0 \pm 11.9^{bc}$<br>(51) | 0<br>(51) |
| | 1-570<br>#23-3 | $4.6 \pm 1.5^c$<br>(57) | $19.7 \pm 15.9^a$<br>(22) | $95.9 \pm 7.4^{bc}$<br>(45) | $4.1 \pm 7.4^{cd}$<br>(45) | 0<br>(45) |
| | 38-570<br>#5-1 | $6.3 \pm 3.4^b$<br>(101) | $20.5 \pm 26.2^a$<br>(66) | $89.3 \pm 9.8^d$<br>(65) | $10.7 \pm 9.8^{ab}$<br>(65) | 0<br>(65) |
| | 38-570<br>#42-1 | $6.0 \pm 1.7^b$<br>(55) | $25.8 \pm 16.4^a$<br>(26) | $94.3 \pm 6.8^c$<br>(42) | $5.8 \pm 6.8^{bc}$<br>(42) | 0<br>(42) |
| Microtubule minus end | SPR2 | NA | $1.5 \pm 2.7^{ef}$<br>(111) | NA | $61.0 \pm 38.0^d$<br>(74) | $39.0 \pm 38.0^a$<br>(74) |
| | *spr2-2* | NA | $7.8 \pm 6.2^b$<br>(68) | NA | $89.3 \pm 20.0^b$<br>(52) | $10.7 \pm 20.0^c$<br>(52) |
| | 1-500<br>#2-2 | NA | $8.6 \pm 6.7^a$<br>(66) | NA | $98.0 \pm 7.3^a$<br>(61) | $2.0 \pm 7.2^d$<br>(61) |
| | 1-570<br>#2-4 | NA | $2.8 \pm 6.3^d$<br>(66) | NA | $77.8 \pm 33.3^c$<br>(59) | $22.2 \pm 33.2^b$<br>(59) |
| | 1-570<br>#23-3 | NA | $4.8 \pm 6.3^c$<br>(60) | NA | $80.4 \pm 27.5^{bc}$<br>(51) | $19.7 \pm 27.5^{bc}$<br>(51) |
| | 38-570<br>#5-1 | NA | $1.2 \pm 2.4^f$<br>(124) | NA | $79.4 \pm 28.4^c$<br>(68) | $20.6 \pm 28.4^b$<br>(68) |
| | 38-570<br>#42-1 | NA | $1.8 \pm 2.2^e$<br>(64) | NA | $84.2 \pm 22.4^{bc}$<br>(38) | $15.8 \pm 22.4^{bc}$<br>(38) |

Values are means ± SD (n = number of events). NA, not applicable. The superscript letters indicate statistically distinguishable groups as determined by the Student's t-test, $p < 0.05$ .

With respect to the cortical microtubule minus ends, both the shortening rate and duration were more severely altered in the 1-500 aa lines compared to the *spr2-2* mutant (Table 1). However, in the 1-570 aa and 38-570 aa lines, these parameters were at least partially restored to control values (Table 1).

## DISCUSSION

The cortical microtubule cytoskeleton of higher plants consists of treadmilling microtubules with free minus ends. A growing body of evidence indicates that minus-end stability is critical for the organization and dynamic remodeling of the cortical microtubule arrays (Fan et al., 2018; Leong et al., 2018; Nakamura et al., 2018). Recently, the SPR2 protein was shown to track microtubule minus ends and inhibit their rapid depolymerization (Fan et al., 2018; Leong et al., 2018; Nakamura et al., 2018). However, the amino-acid sequence of SPR2 is unrelated to the CAMSAP/Patronin/Nezha family of −TIPs, raising the question of how SPR2 recognizes and stabilizes microtubule minus ends. Here, we used a combination of biochemical, cell biological and genetic analyses to elucidate the structural determinants of microtubule and tubulin binding, and of minus-end stabilization by SPR2.

We demonstrate that the N-terminal region of SPR2 contains an active TOG domain that binds to soluble tubulin. Based on our structure-function analysis, we predict that SPR2 contains a sixth HEAT repeat between residues Ala38 and Ile70. Importantly, multimerization via the coiled-coil domain is vital for SPR2 to bind to tubulin. TOG domain-containing microtubule regulatory proteins that do not contain a linked array of TOG domains frequently multimerize (Farmer and Zanic, 2021). For example, Cep104 and CLASP, which contain one and two TOG domains respectively, possess a coiled-coil domain that mediates homodimerization (Al-Bassam et al., 2010; Al-Jassar et al., 2017). Analogous to SPR2, homodimerization of CLASP is required to bind tightly to tubulin dimers (Al-Bassam et al., 2010).

Similar to XMAP215 and CLASP (Al-Bassam et al., 2010; Widlund et al., 2011), SPR2 contains a basic region that contributes to microtubule lattice binding. However, the basic region of SPR2 is not sufficient for lattice binding, even when paired with the coiled-coil domain. In addition, deleting the first 70 aa of SPR2 abrogates microtubule binding. Since the 1-276 aa fragment is not sufficient for microtubule binding, we conclude that the 1-70 aa region is necessary but not sufficient for SPR2 to bind to microtubules. In any case, multimerization of SPR2 greatly enhances microtubule binding, likely due to multivalent interactions with the microtubule lattice. Microtubule-bound multimers of SPR2 can also bind to soluble tubulin, indicating that lattice and soluble tubulin binding are separable functions.

The 1-400 aa fragment of SPR2 binds to the microtubule lattice while the 1-500 aa fragment binds to both the microtubule lattice and minus ends. These data indicate that the 115 aa segment between the basic region and coiled-coil domain somehow confers minus end localization. This region is predicted to be disordered and it is possible that it mediates recognition of a specific tubulin conformation found at the minus end.

Genetic complementation tests showed that expression of (1-500)-mRuby is not sufficient to rescue the helical growth phenotype of the Arabidopsis *spr2-2* mutant. Unlike full-length SPR2 which continuously tracks depolymerizing cortical microtubule minus ends, (1-500)-mRuby labels depolymerizing minus ends sporadically. In addition, the 1-500 aa fragment lacks tubulin binding ability *in vitro*. Taken together, these data suggest that tubulin binding and stable minus end tracking are crucial to the function of SPR2. By contrast, the (1-570)-mRuby expressing plants showed either partial or complete complementation. We found that the fluorescence signal of the (1-570)-mRuby fusion protein is higher in plants that show complete complementation than in plants that exhibit partial complementation. However, we were unable to compare protein levels more rigorously between these plants because of the lack of suitable antibodies for SPR2 and mRuby. Importantly, the 38-570 aa fragment showed complete complementation when expressed in the *spr2-2* mutant. This fragment best recapitulates the microtubule binding and minus end stabilizing activity of full-length SPR2 *in vitro*. Likewise, the (38-570)-mRuby fusion protein showed continuous tracking of depolymerizing cortical microtubule minus ends *in planta*. Thus, while the 1-37 aa Ser/Thr-rich region is necessary for SPR2 to track microtubule minus ends *in vitro*, it is not necessary for this ability *in planta*.

Our work demonstrates that the C-terminal domain is not vital to the microtubule minus-end stabilizing function of SPR2. Nonetheless, it is important to note that the dynamics of cortical microtubules are not fully rescued in the *spr2-2* mutant expressing either (1-570)-mRuby or (38-570)-mRuby. Future experiments will investigate whether and how the C-terminal domain contributes to SPR2 function. Since the C-terminus is predicted to contain armadillo repeats, one possibility is that it engages in protein-protein interactions that regulate the localization and/or activity of SPR2. Another possibility, based on the fact that SPR2 does not autonomously target microtubule plus ends (Fan et al., 2018), is that the C-terminus interacts with +TIPs to robustly target SPR2 to the growing microtubule plus end. Our finding that the (38-570)-mRuby fusion protein decorates growing cortical microtubule plus ends less frequently than full-length SPR2-mRuby supports the latter possibility.

How might SPR2 inhibit the depolymerization rate of the microtubule minus end? Based on our data, we propose that SPR2 stably associated with the microtubule minus end captures soluble tubulin. During depolymerization of the minus end, SPR2 lets go of its bound tubulin as it tracks the shortening end. This activity would locally increase tubulin concentration and slow down the depolymerization rate. Regardless of the exact mechanism, our data establishes SPR2 as the first microtubule regulatory protein that uses a TOG domain to regulate the dynamics of microtubule minus ends. XMAP215 homologs were recently found to also localize to microtubule minus-ends (Gunzelmann et al., 2018; Thawani et al., 2018). However, their TOG domains do not stabilize minus-ends but rather promote nucleation by facilitating the binding of tubulin dimers to the γ-tubulin ring complex (Gunzelmann et al., 2018; Thawani et al., 2018). Therefore, the SPR2 protein provides a unique opportunity to study how nature has co-opted TOG domains for microtubule minus-end stabilization.

## MATERIALS AND METHODS

### Plant growth

*Arabidopsis thaliana* (L.) Heynh. Columbia-0 (Col-0) plants were used for all experiments. For growth on plates, seeds were sterilized by 25% (v/v) bleach for 10 min, rinsed five times with sterile ultrapure water, suspended in 0.1% (w/v) sterile agarose solution and planted on plates containing 2.2 g/L MS salts with Gamborg’s vitamins (Caisson Laboratories), 3 g/L Sucrose, pH 5.7. Seeds were stratified in the dark at 4°C for 2-3 days and then grown at 23°C under 16h light for 4 days. When transferred to the soil, seedlings were grown under continuous light at 22°C.

### Constructs

For live imaging of SPR2 fragments, the *pSPR2∷SPR2-mRuby* construct (Fan et al., 2018) was modified by replacing the full-length *SPR2* cDNA with the appropriate truncated *SPR2* cDNAs. These constructs were introduced into the *spr2-2* mutant expressing a GFP-TUB6 microtubule marker (Fan et al., 2018) using *Agrobacterium*-mediated floral dip transformation. Transgenic plants were selected using 100 mg/L gentamicin with 50 mg/L kanamycin, and homozygous lines containing a single copy of the transgene were used for study.

For recombinant protein expression, the pGEX *SPR2-GFP* construct (Fan et al., 2018) was modified by inserting a PaeI restriction site between *SPR2* and *GFP* and the PreScission protease site was replaced by a TEV protease site. *SPR2* fragments were generated using PCR, restriction digested by BamHI and PaeI enzymes and used to replace the full-length *SPR2* in the modified pGEX *SPR2-GFP* construct. Constructs were introduced into *E. coli* Rosetta (DE3) cells for protein expression. Primers used in this study are listed in Supplemental Table 1.

### Protein purification

GST-tagged proteins were purified using affinity chromatography with Glutathione Sepharose beads (GE Healthcare) in a cold room. Purified proteins were subsequently treated with His-tagged TEV protease (1.7 mg/ml) in cleavage buffer (50 mM Tris-HCl, 150 mM NaCl, 1 mM EDTA, 1 mM DTT, 1% Triton X-100, pH 7.0) for 20 hours in a cold room to remove the GST tag, and free SPR2 proteins were separated from the beads by centrifugation. The TEV protease was removed from the digested proteins by incubation with Ni-NTA beads for 2.5 hours in a cold room. Isolated proteins were desalted using PD-10 columns (GE Healthcare) and exchanged into BRB80 buffer (80 mM piperazine-1,4-bis(2-ethanesulfonic acid), 1mM MgCl2, and 1mM EGTA, pH 6.8) supplemented with 50 mM NaCl to prevent protein aggregation.

### Microtubule cosedimentation

Microtubules were prepared using unlabeled porcine brain tubulin (Cytoskeleton) reconstituted in ice-cold BRB80 buffer to 5 mg/ml and polymerized in the presence of 1mM Mg-GTP for 1 hour at 37°C. The assembled microtubules were stabilized with 20 μM paclitaxel (Cytoskeleton) and stored at room temperature. For microtubule cosedimentation experiments, 2 μM SPR2(1-500)-GFP or SPR2(1-570)-GFP were coincubated with different concentrations of taxol-stabilized microtubules, 0.1 mg/ml bovine serum albulmin, and 20 μM paclitaxel for 20 min. Microtubules and bound proteins were sedimented by centrifugation at 100,000 g at 25°C for 20 min. Supernatant and pellet fractions were resuspended in equal volumes of loading buffer and analyzed using SDS-PAGE. The amount of SPR2 protein in the supernatant and pellet fractions was quantified using densitometry.

### *In vitro* functional reconstitution experiments

*In vitro* reconstitution assays with dynamic microtubules were conducted in flow chambers assembled using glass slides and silanized coverslips attached with two pieces of double-sided sticky tape. The flow chambers were first coated with 20% monoclonal anti-Biotin antibody (Sigma-Aldrich) in BRB80 and then blocked by 5% Pluronic F-127 (Sigma-Aldrich) in BRB80 for 5 min each. GMPCPP-stabilized microtubule seeds were assembled using 50 μM porcine tubulin containing 1:12 biotin-labeled and 1:10 rhodamine-labeled porcine tubulins (Cytoskeleton) and polymerized in the presence of 1mM GMPCPP (Jena Bioscience) at 37°C for 30 min. The polymerized seeds were then fragmented by passing through a 100 mL Hamilton syringe five times. 300 nM GMPCPP-stabilized seeds were introduced into the flow chamber and allowed to bind to the anti-biotin antibody for 10 min. Microtubule polymerization was initiated by flowing in 20 mM 1:25 rhodamine-labeled porcine tubulin in BRB80 buffer containing 1% methyl cellulose (4000 cP, Sigma-Aldrich), 50 mM DTT, 2 mM GTP and an oxygen-scavenging system consisting of 250 mg/ml glucose oxidase, 35 mg/ml catalase and 4.5 mg/ml glucose. 500 nM of full-length or truncated SPR2-GFP proteins were included in the polymerization mix to study their microtubule localization and effect on microtubule dynamics. GFP and rhodamine were excited using 5-mW 488-nm and 561-nm diode-pumped solid-state lasers, respectively. Images were collected by a 100X (NA 1.45) objective and a back-illuminated electron-multiplying CCD camera (ImageEM; Hamamatsu) at 2 s intervals.

### Live imaging

Live imaging was conducted using variable-angle epifluorescence microscopy. Images were collected from epidermal cells in the apical hypocotyl region of four-day-old, light-grown seedlings. Seedlings were gently mounted in water between two layers of double-sided adhesive tape on a slide and covered with No. 1.5 glass coverslip. GFP was excited using 2-mW, 488-nm diode-pumped solid-state laser (Melles Griot), and mRuby was excited using 2-mW 561-nm diode-pumped solid-state laser (Melles Griot), respectively. Images were collected by a 100X (NA 1.45) objective and a back-illuminated electron-multiplying CCD camera (ImageEM; Hamamatsu) at 2 s intervals.

### Soluble tubulin pulldown

Anti-GFP beads (MBL Life Science) were washed twice with binding buffer (BRB80 buffer supplemented with 50 mM NaCl and 0.05% Tween-20). The equilibrated beads were incubated with 1 μM full-length SPR2-GFP protein on a rotary shaker for 2.5 hours at room temperature. Then, 3 μM BSA was added and incubated for an additional 1 hour on a shaker in a cold room. Subsequently, 2 μM unlabeled porcine tubulin (Cytoskeleton) prepared in binding buffer containing 10 mM DTT and 0.2 mM PMSF was added and incubated overnight on a shaker in a cold room. Beads were then collected by centrifugation at 1000 g for 1 min and washed 3-times with binding buffer. Beads were transferred to fresh tubes and washed for an additional 5-times with binding buffer. Bound proteins were then analyzed with SDS-PAGE. As controls, 1 μM of full-length SPR2-GFP and 2 μM unlabeled porcine tubulin were incubated separately with the anti-GFP beads.

### Imaging soluble tubulin binding to microtubule-bound SPR2-GFP proteins

GMPCPP-stabilized microtubules were assembled using 50 μM unlabeled porcine tubulin containing 1:12 biotin-labeled porcine tubulin (Cytoskeleton) and polymerized in the presence of 1mM GMPCPP (Jena Bioscience) at 37°C for 30 min. 300 nM GMPCPP-stabilized microtubules were introduced into a flow chamber and allowed to bind to anti-biotin antibody for 10 min. 100 nM of the specified SPR2-GFP proteins along with 2.5 μM 1:9 rhodamine-labeled porcine tubulin were then introduced into the flow chamber in BRB80 buffer containing 1% methyl cellulose (4000 cP, Sigma-Aldrin), 50 mM DTT and an oxygen-scavenging system consisting of 250 μg/ml glucose oxidase, 5 μg/ml catalase and 4.5 mg/ml glucose. GFP and rhodamine were excited using 5-mW 488-nm and 561-nm diode-pumped solid-state lasers, respectively. Images were collected by a 100x (NA 1.45) objective and a back-illuminated electron-multiplying CCD camera (Image EM; Hamamatsu) at 2 s intervals.

### Native PAGE electrophoresis

The multimerization status of full-length and specified truncated SPR2-GFP proteins was analyzed using the NativePAGE Bis-Tris gel system (ThermoFisher Scientific). 2 μM of purified proteins were loaded onto a 4-16% NativePAGE gel and ran according to the manufacturer’s procedure at 150 V constant for about 120 min.

### Quantification and statistical analysis

All experiments were repeated at least three times using independent plants and protein preparations. Statistical significance of data was calculated using the Student’s *t* test. To quantify the dynamics of microtubules *in vivo* and *in vitro*, kymographs of individual microtubules were generated using the Image J/Fiji Dynamic Kymograph plugin (Zhou et al., 2020). Microtubule growth and shortening rates, and the duration of growth, shortening, and pause were calculated using the Microtubule Kymograph Analysis plugin (Zhou et al., 2020). Graphs and curve fitting was done using GraphPad Prism.

## ACKNOWLEDGMENTS

This work was supported by the National Institute of General Medical Sciences of the National Institutes of Health under award number R35GM139552. N.B. was supported by a William H. Danforth fellowship in plant sciences.

---

**Supplemental Table 1:**
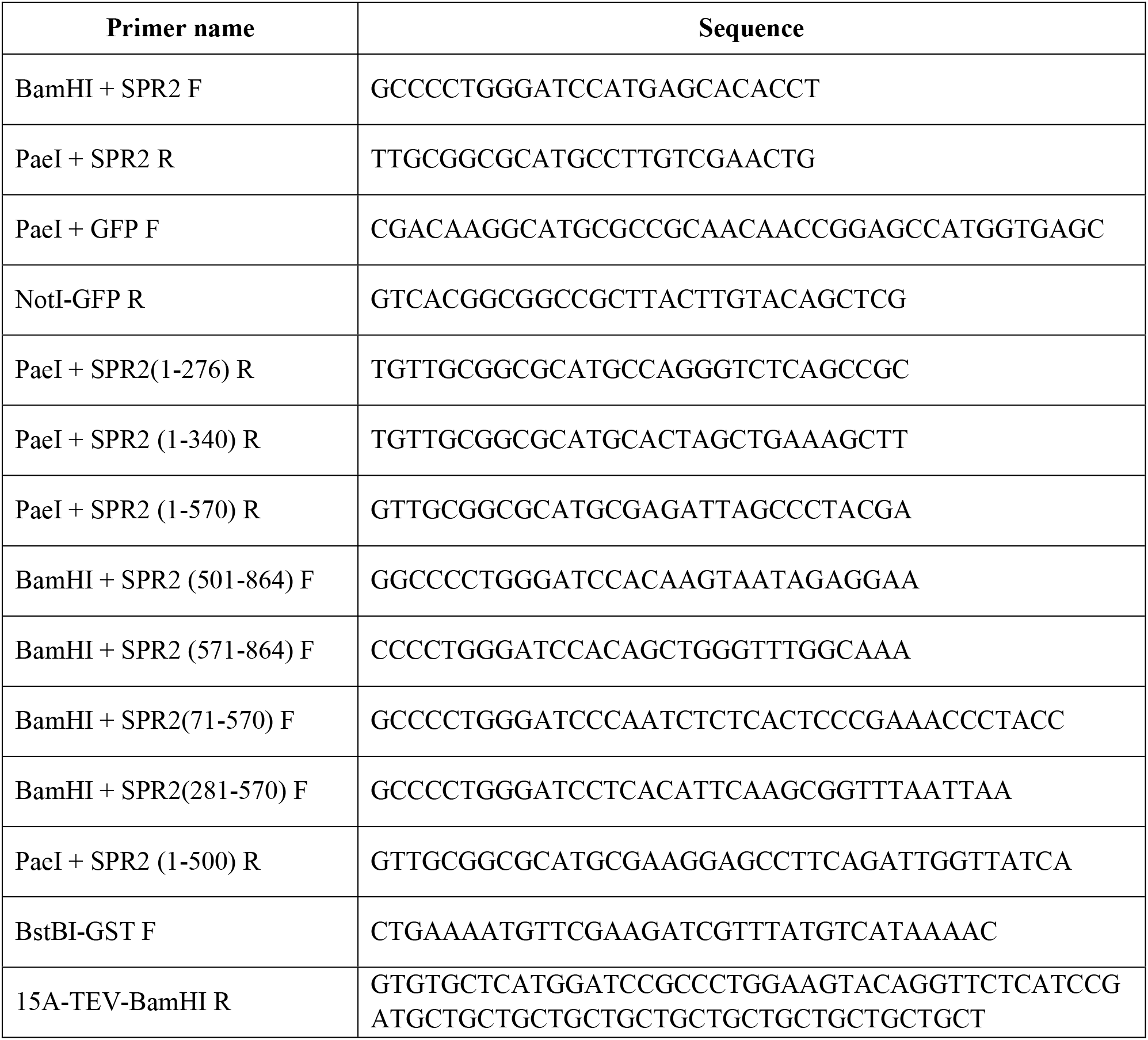
Oligonucleotides used in this study

